# Circadian IOP rhythm in rats is driven by neural signals from the brain

**DOI:** 10.1101/2025.10.27.684436

**Authors:** Alexandra Zamitalo, Christopher L. Passaglia

**Affiliations:** Medical Engineering Department, University of South Florida, Tampa, FL 33620; Ophthalmology Department, University of South Florida, Tampa, FL 33620

## Abstract

Intraocular pressure (IOP) exhibits a robust circadian rhythm whose driving mechanisms remain poorly defined. This study aimed to establish whether the circadian IOP rhythm in rats originates from local ocular oscillators or non-local neural or humoral inputs. IOP was monitored continuously in adult Brown Norway rats using an in-house telemetry system, and topical neurotoxin instillation and superior cervical ganglionectomy were performed. TTX produced a rapid, dose-dependent reduction of nocturnal IOP when applied during the dark phase but had no effect during the light phase. This hypotensive effect was unilateral, short-lived, and consistent with local sodium channel blockade of sympathetic efferents. In contrast, SCGx abolished the circadian rhythm entirely, eliminating the nocturnal IOP elevation. These findings demonstrate that the rat IOP rhythm is not generated by an intrinsic oscillator or circulating humoral signals but is instead driven by sympathetic efferent input from the superior cervical ganglion. These results highlight species-specific differences in the mechanisms of circadian IOP regulation and motivate future direct mechanistic studies on the IOP rhythm in relevant animal models of glaucoma.

## INTRODUCTION

Glaucoma is the leading cause of irreversible vision loss worldwide, and intraocular pressure (IOP) is a key parameter for detecting and managing disease onset and progression [1-4]. Changes in mean IOP often garner most clinical and research attention, but other aspects of IOP variation may contribute significantly to glaucoma pathophysiology as well. This is because IOP is highly dynamic, fluctuating over timescales of seconds to days due to a variety of internal and external factors. Amongst the largest factors is the diurnal IOP rhythm that is expressed by many mammalian species [5-9], including humans [10, 11]. The rhythm causes IOP to peak during the night and trough during the day, even under constant lighting conditions [7, 46-49]. Disruption of this rhythm has been correlated with optic nerve damage, suggesting a possible role in glaucoma development [10, 12-17]. It is thereby of paramount importance to understand the origin and function of circadian IOP rhythmicity.

A fundamental question is whether the IOP rhythm is driven by a central or peripheral clock. Circadian rhythms are synchronized throughout the body by a network of clock neurons in the hypothalamus of the brain, which can entrain to ambient light cycles via visual input from the eye. The central clock sends circadian messages through neural and hormonal pathways that can directly generate rhythmicity in peripheral tissues or modify endogenous rhythms generated by local clocks in those tissues. Peripheral clocks have been implicated in liver metabolism [18-21], glucocorticoid production in the adrenal glands [22-24], and barrier functions in the lungs [25, 26] and skin [27, 28]. They have also been identified in several ocular tissues, including the retina [29-32], cornea [33-35], and iris-ciliary body [36-38]. The retinal clock is known to drive a melatonin rhythm that alters retinal gene expression, circuit connectivity, and light sensitivity during the day and night [39-43]. None of the ocular clocks have been shown to drive the IOP rhythm as yet [10, 44, 45].

The mechanisms of IOP rhythmogenesis may differ across animals. In mice, peripheral clocks in the iris-ciliary body complex can influence the phase and amplitude of the IOP rhythm but cannot support the rhythm [36, 50]. Non-local neural and hormonal pathways are required that involve circadian sympathetic nerve transmission and glucocorticoid/corticosterone circulation to mice eyes [44, 45, 51]. In rabbits, on the other hand, sympathetic nerve signals from the central clock increase IOP at night [8, 52, 53] and another unknown process lowers IOP during the day [54]. The objective of this work was to elucidate the origin of circadian IOP rhythmicity in rats. Rats are a popular animal model for glaucoma research, along with mice and rabbits, and they may provide valuable insights about circadian clock contributions to the disease in humans.

## METHODS

All experiments were conducted on adult Brown-Norway rats (male, retired breeder, 300-400g) in compliance with the National Institutes of Health guide for the care and use of laboratory animals, ARVO Statement for the use of animals in ophthalmic and vision research, and protocols approved by the Institutional Animal Care and Use Committee at the University of South Florida. Animals were housed prior to experimentation under a 12-hr light (6AM-6PM) / 12-hr dark (6PM-6AM) cycle in a humidity- and temperature-controlled room with food and water freely available. Animals were then outfitted with an in-house wireless telemetry system that continuously recorded IOP round-the-clock at 0.25 Hz. Details of the IOP telemetry system have been published [55, 56]. Briefly, a drug-loaded silicone microcannula was surgically implanted in the anterior chamber of one or both eyes, secured to the sclera with sutures, and routed subdermally to a skull-mounted coupler that was connected fluidically to a pressure sensor worn on the animal’s back. After cannulation surgery, animals were returned to housing for at least 3 days to record baseline IOP rhythmicity under the ambient light/dark (LD) cycle and then subjected to testing.

### Neurotoxin instillation

A cohort of unilaterally-canulated animals (n=8) and a cohort of bilaterally-cannulated animals (n=4) animals were transferred to an environmental control unit (ECU, BIO-C36; Tecniplast, Buguggiate, Italy) that allowed for manipulation of ambient lighting. After 3 days of LD entrainment, animals were placed in constant darkness (DD) to eliminate any direct effects of light on IOP. A 10uL drop of saline or tetrodotoxin (TTX), a voltage gated sodium channel blocker, was then instilled on the cannulated eye at different times of day and concentrations (0.001, 0.1, 0.5, 1 mg/mL). The drop was applied to only one eye of bilaterally-cannulated animals. Instillations were performed under dim red light and brief (<3 min) 2% isoflurane at least 24-hrs apart. They were administered at distinct phases of the free-running IOP rhythm in DD, with trough and peak phases corresponding in circadian time (CT) to subjective day (SD) and subjective night (SN) when light would be respectively on and off. SD thereby spanned from CT0-12 and SN from CT12-24.

### Superior cervical ganglionectomy

A separate cohort of unilaterally-cannulated animals (n=3) underwent a bilateral superior cervical ganglionectomy (SCGx). The animals were administered extended-release analgesics and anesthetized via intraperitoneal injection of ketamine (1mg/mL) and xylazine (0.1mg/mL). The nape of the neck was shaved, disinfected, and a vertical incision (∼25mm) made to expose the mandibular glands. The glands were slid aside to access the carotid triangle, and the carotid arteries were revealed by blunt dissection. A 4-0 suture was fed under the arteries to gently retract the vessels without completely obstructing blood flow. The underlying superior cervical ganglion (SCG) was removed by transecting the sympathetic trunk and carotid nerves on the distal and proximal sides of the ganglion. The procedure was repeated on the contralateral SCG, the neck incision was closed with resorbable sutures, and topical antibiotics were applied daily for 2-3 days. SCGx success was confirmed qualitatively by monitoring eyelid ptosis [57]. Animals that did not exhibit signs of SCGx success were omitted from the study. Following surgery, animals were returned to housing and IOP was monitored under the ambient LD cycle until experiment termination.

### Data Analysis

IOP data were processed in MATLAB (The Mathworks, Natick, MA, USA) using median and mean filters of 28-second width to remove outliers and smooth records. The IOP rhythm in DD can drift out of synch with the external LD cycle so the amplitude, phase, and period of the free-running rhythm and TTX-induced changes in these parameters were estimated by cosinor analysis [46]. Specifically, IOP records of at least 36-hr duration before and after TTX instillation were respectively fit to a single cosinor function:

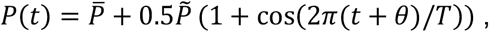

where 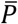 is baseline IOP during SD, 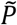 is peak-to-trough rhythm amplitude, *T* is rhythm period, and *θ* is rhythm phase with respect to midnight. The magnitude of TTX effects was quantified by comparing mean IOP for the 2-hr interval immediately before and after instillation. The duration of TTX effects was defined as the time from instillation until IOP recovery to pre-instillation level. The effect of TTX concentration was quantified by fitting IOP data over a 24-hr period with a difference-of-pulses function given by:

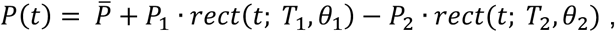

where 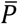 is baseline IOP during SD, *P*_1_ and *P*_2_ are the amplitude of the two pulses, *T*_1_ and *T*_2_ are the pulse durations, and *θ*_1_ and *θ*_2_ are pulse phases with respect to instillation time. Fits were obtained by nonlinear regression using criteria of *T*_1_ > *T*_2_ and *θ*_2_ > 0 > *θ*_1_. SCGx effects were quantified by comparing mean IOP over a 4-hr interval during SD (CT4-CT8) and SN (CT16-CT20) for the 3 days before and after the surgical procedure. Statistical analyses were performed with SigmaPlot software (Systat, San Jose, CA, USA) using paired Student’s t-test or one-way repeated measures ANOVA with Holm-Sidak method for multiple comparisons on normally distributed data and Mann-Whitney rank sum tests on non-normal data. Results are reported as mean ± standard deviation or median [upper quartile, lower quartile] for normal and non-normal data, respectively. Significance was defined as p<0.05.

## RESULTS

Data were collected from a total of 14 rats. All animals exhibited a pronounced IOP rhythm that was synchronized to the ambient LD cycle. Fig 1A shows that the rhythm persisted in DD as well. TTX was applied topically at different times of day and night to block voltage-gated sodium channels and determine if the IOP rhythm depends on nerve impulses. Fig 1B shows 2-day IOP records during which TTX was instilled at CT3, CT11, CT15, and CT19, which corresponded to around 9AM, 5PM, 9PM, and 1AM on a 24-hour clock. HERE During SN, TTX temporarily reduced IOP for several hours, at CT11, it delayed nocturnal IOP elevation compared to the previous night, and during SD it had no effect. Fig 1C demonstrates that SN saline instillation under identical conditions had no effect on IOP (21.7 ± 7.0mmHg to 21.5 ± 5.7mmHg, n=4, p=0.82). Fig 1D summarizes differences in IOP response to TTX across the 6 times tested. TTX did not alter IOP at CT3 (p=0.13), CT7 (p=0.24), or CT11 (p=0.10) but significantly reduced it at CT15 (20.2 ± 3.6mmHg to 11.3 ± 2.5mmHg, p=0.002), CT19 (19.4 ± 3.9 mmHg to 10.4 ± 2.2mmHg, p=0.01), and CT24 (13.7 ± 2.0mmHg to 12.3 ± 1.4mmHg, p=0.04). This time-dependent IOP response to TTX is consistent with a 24-hr baseline IOP not under neural control that is elevated during SN by neural input to the eye.

**FIGURE 1.**
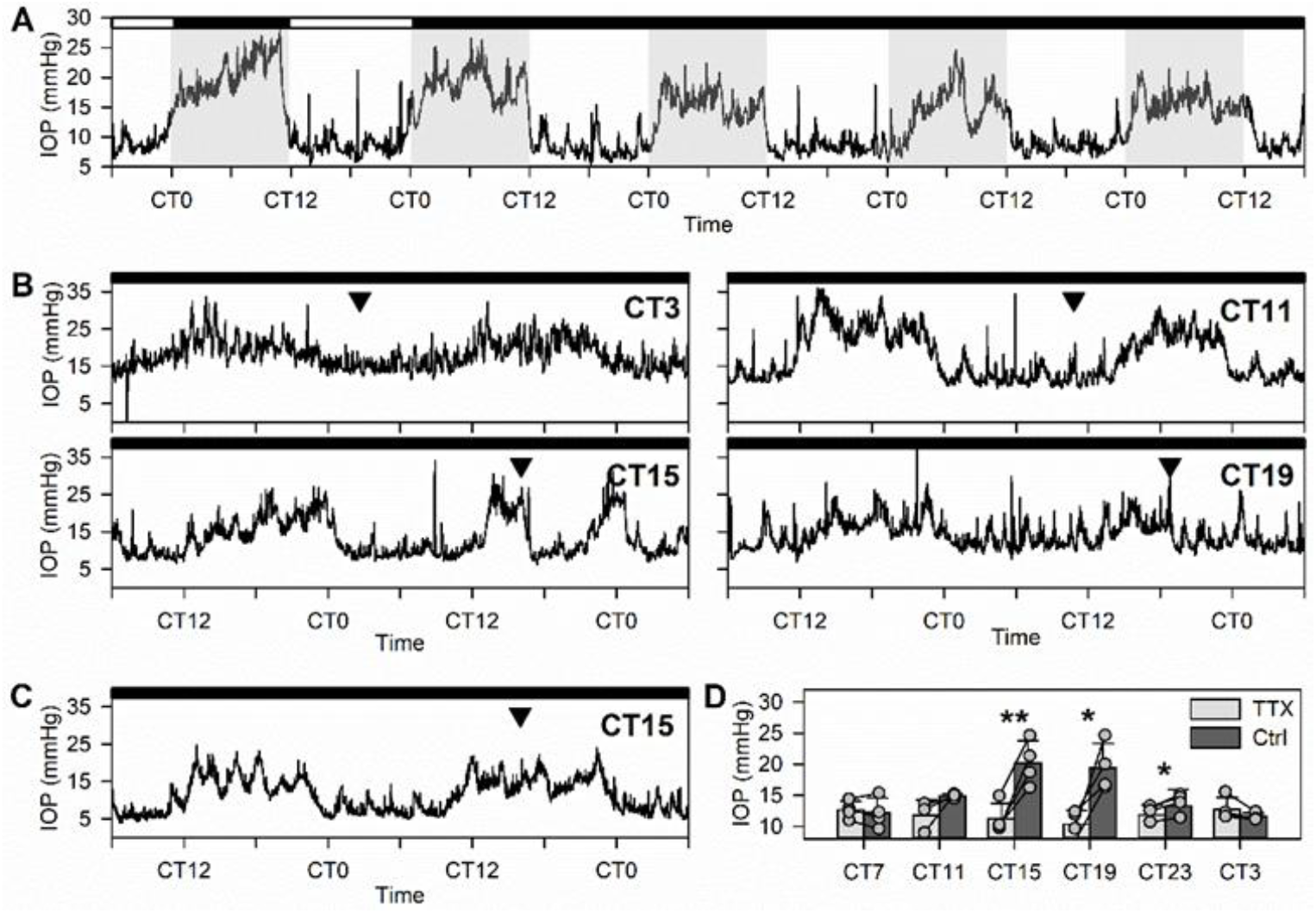
Time dependent TTX effect on IOP. A: The circadian IOP rhythm entrains to ambient light and persists in the absence of photic entrainment cues. Grey shading represents the dark phase of the LD cycle to which the animal was entrained. Black and white bars (top) represent times when the lights were off and on in the ECU, respectively. B: Representative IOP responses to TTX at various instillation times. Instillation is indicated by the black arrowhead. C: Representative IOP response to saline instilled at the time indicated by the black arrowhead. D: Mean IOP comparison before and after TTX instillation at different times. Light bars represent post-TTX IOP and dark bars represent pre-TTX IOP. ^*^ = p<0.05, ^**^ = p<0.01.

The dose dependence of the IOP response to TTX was assessed to deduce whether SN IOP could be reduced only partially to daytime levels (n=3). Fig 2 shows 24-hr IOP records fit with a sum of two pulses to model the behaviors of the circadian rhythm and the TTX response. The amplitude of the maximal TTX effect was represented as a percentage of the nocturnal IOP elevation and exhibited a graded response proportional to TTX concentration. 1mg/mL TTX reduced IOP by 103% ± 15% of the circadian rhythm amplitude, 0.5mg/mL by 84% ± 24%, 0.1mg/mL by 77% ± 13%, and 0.01mg/mL by 38% ± 17%. Significant differences were observed between 1mg/mL and 0.1mg/mL (p=0.02), 0.1mg/mL and 0.01mg/mL (p=0.05), and 1mg/mL vs 0.01mg/mL (p=0.03) but not between 1mg/mL and 0.5mg/mL (p=0.14), 0.5mg/mL and 0.1mg/mL (p=0.40), or 0.5mg/mL and 0.01mg/mL (p= 0.09). This graded effect is likely due to TTX blocking more nerve projections to the anterior segment at higher doses.

**FIGURE 2.**
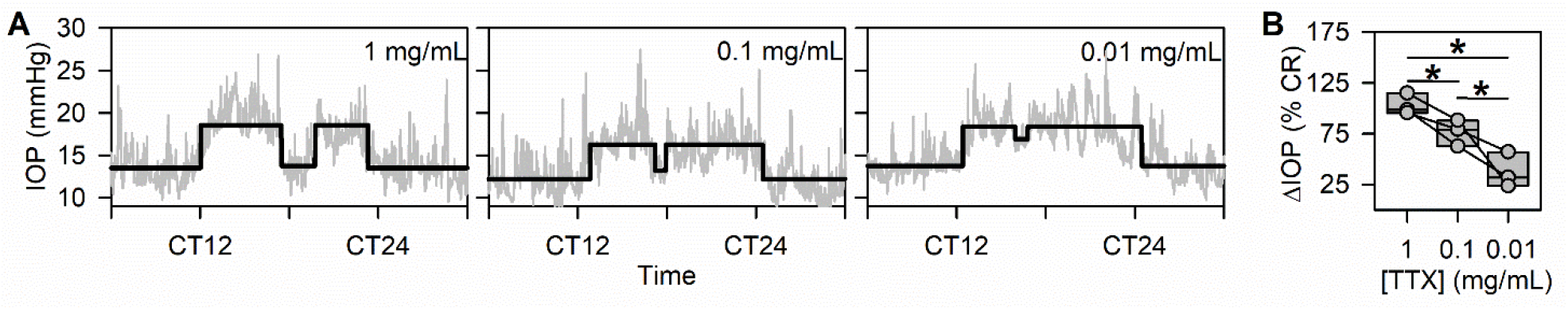
TTX affects SN IOP in a dose-dependent manner. A: Representative IOP responses at the various doses tested (grey) with the output of the fit describing the TTX effects overlaid (black). B: Comparison of TTX effect magnitude by dose as a percentage of the amplitude of the nocturnal IOP elevation. ^*^ = p<0.05.

The time course of the IOP response to TTX was assessed using the CT18 experiments (n=4). Fig 3A shows a 24-hr IOP record in which TTX was instilled during SN, and the duration of the effect is labelled with a bar. TTX induced IOP hypotension lasted 4.7 ± 1.5 hours. Fig 3B summarizes the onset dynamics of the IOP response. The half-maximal and maximal effects occurred 16 ± 8.1 mmHg and 93 ± 36 minutes after instillation, corresponding to 6.0 ± 2.9% and 35 ± 17% of the overall response duration, respectively. The hypotensive effect occurs on the order of tens-of-minutes exhibiting little to no lag after instillation which is consistent with a local neural effect rather than a systemic one.

**FIGURE 3:**
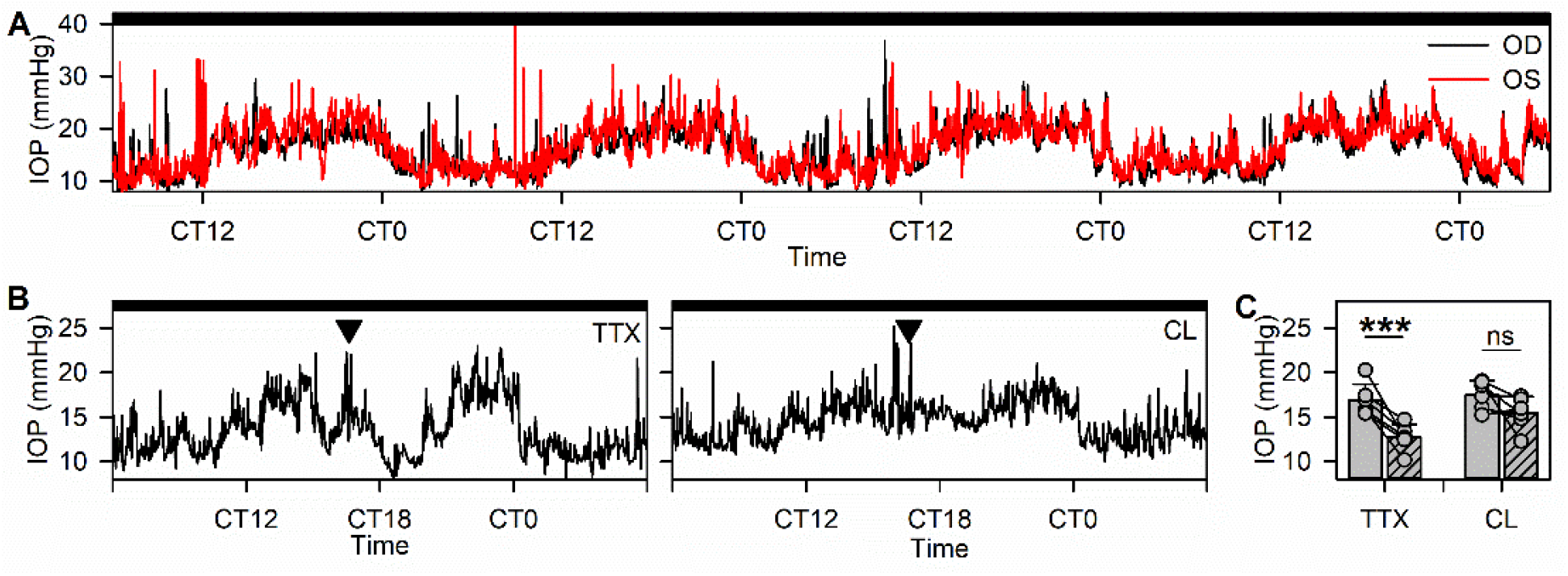
TTX effects are limited to the eye on which it was applied. A: Representative IOP records from paired eyes. Red = OS, black = OD. B: Paired IOP records during which TTX was applied to one eye (left) and effects were studied in the CL eye (right). C: Statistical comparison of pre- and post-TTX IOP in the treatment (TTX) and contralateral (CL) eye. Light bars represent the pre-instillation IOP, and dark bars represent post-instillation IOPs. ^***^ = p<0.005, ns = not significant.

Fig 4A shows 4-day IOP records from paired eyes demonstrating that circadian behavior is generally mirrored in paired eyes, consistent with synchronization by a central master clock. TTX was applied topically to only one eye of bilaterally cannulated animals (n=4) to determine if the observed effect was the result of systemic uptake of the drug from the corneal surface. Fig 4B shows paired 24-hour IOP records demonstrating that only the experimental eye responded to TTX instillation. Fig 4C summarizes this observation across n=6 drops showing that the experimental IOP was reduced from 16.9 ± 1.8 mmHg to 12.7 ± 1.5 mmHg (p=0.002) while contralateral IOP was not changed (p=0.11). Because no response was observed in the untreated eye, the TTX effect must not be prompted by a systemic change.

**FIGURE 4:**
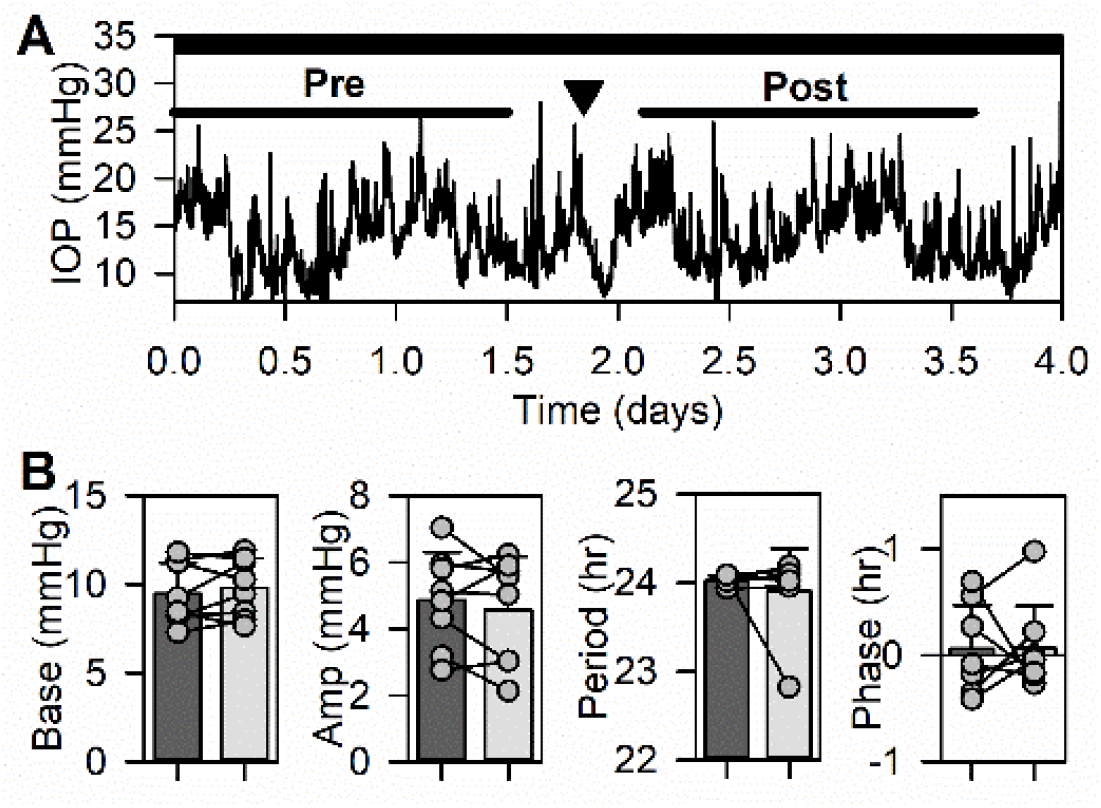
TTX did not alter circadian rhythmic behavior of the following day. A: 4-day IOP record annotated to show time of TTX instillation (black arrowhead) and data segments used for cosine fitting pre- and post-instillation. B: Summary of TTX effects on IOP baseline (Base), circadian amplitude (Amp) and rhythm period and phase. Dark bars represent pre-instillation data; light bars represent post-instillation data. None of the data displayed were significantly different.

A cosine waveform was fit to at least 36 hr IOP records prior to and following TTX instillation to identify whether TTX was acting on voltage-gated sodium channel-dependent processes in the retina including SCN entrainment. Fig 5A defines which IOP segments were used for cosinor analysis. Fig 4B summarizes rhythm parameters before and after TTX across animals (n=4). The IOP rhythm baseline, amplitude, period, and phase were unchanged (p=0.20, p=0.20, p=0.38, and p=0.74, respectively) which shows that TTX instillation does not affect the free-running IOP rhythm and thus must be acting only on efferent synchronization not on entrainment.

**FIGURE 5:**
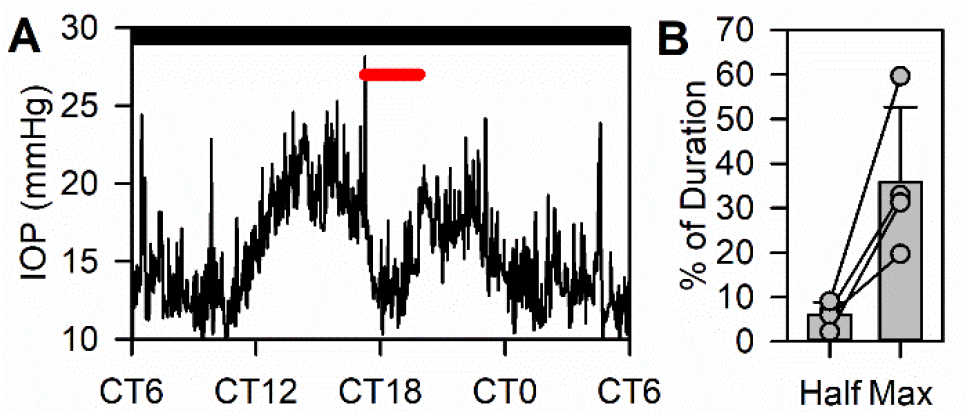
IOP response to SN TTX instillation is rapid. A: Representative 24-hour IOP record with the TTX effect duration labelled in red. B: Comparison of time to half effect (Half) and max effect (Max) represented as a percentage of total effect duration.

SCGx was performed on a separate cohort by removing the bulbous structure positioned behind the carotid bifurcation shown in Fig 6A. IOP was recorded in 3 rats for 6.4 ± 3.8 days after SCGx. Fig 6B shows a representative 5-day IOP record exhibiting a clear and pronounced circadian rhythm that was knocked out by SCG removal. Mean IOP during SD and SN were compared to detect circadian variation in IOP. Fig 6C demonstrates that IOP during SD (11.1 ± 1.6 mmHg) was not different from SN (12 ± 1.7 mmHg, p=0.1) indicating that the IOP rhythm was abolished. This finding confirms that the circadian IOP rhythm is not generated by a local oscillator within the ocular tissue but instead requires neural input.

**FIGURE 6:**
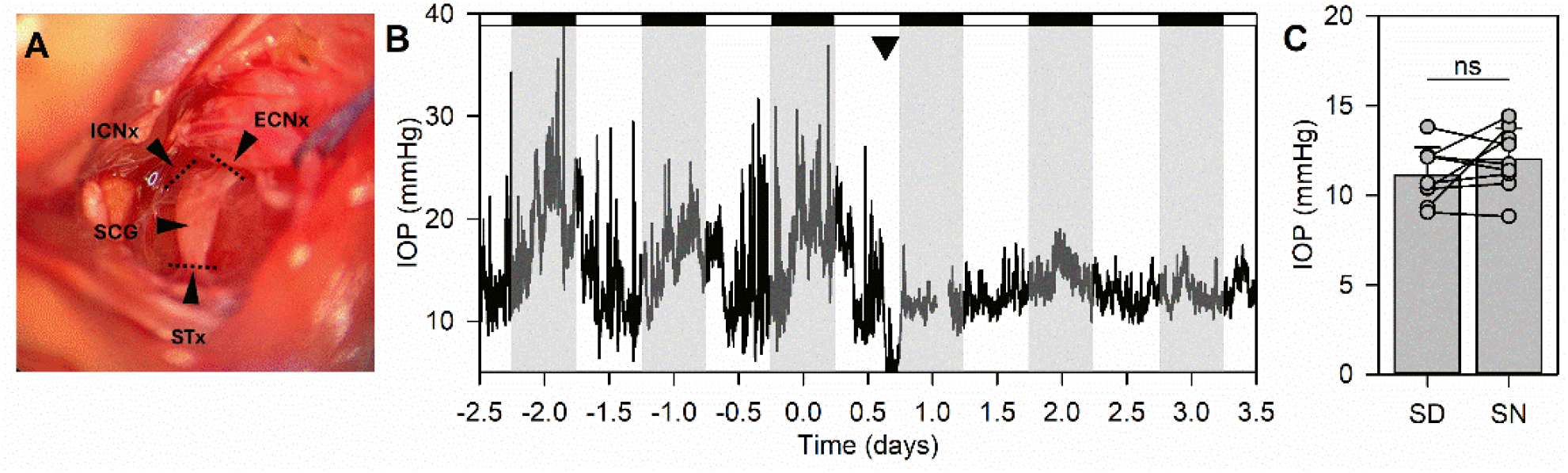
SCGx abolishes the IOP rhythm. A: Image of SCG with incision sites labelled with dotted lines through the major projections off the ganglion. B: Representative 6-day IOP record during which SCGx was performed (black arrowhead). Grey shading represents dark phases of the entrainment cycle. White and black bars (top)represent lights on and off in the ECU, respectively. C: IOP comparison during SD and SN in the days following SCGx. ns = not significant.

## DISCUSSION

### Summary of Results

In this study, the circadian IOP was disrupted with various approaches targeting sympathetic input to the eye. The effects of these techniques on IOP were captured in state-of-the-art temporal resolution in ambulatory animals, offering previously unattainable insight into the timing of these responses. As shown previously with both tonometry and telemetry-based studies, IOP exhibited a strong diurnal rhythm with peak values occurring in the dark phase and troughs during the light phase. The nocturnal IOP elevation in rats appears to be driven by the sympathetic nervous system as opposed to a local ocular oscillator or changes in circulation of signaling molecules. This is supported primarily by two findings. First, topical TTX instillation at several times in the circadian cycle revealed that nocturnal IOP requires voltage gated sodium channel activation, whereas daytime IOP does not. Second, sympathetic silencing by SCGx completely abolished the IOP rhythm in subsequent days. These results reveal a stronger dependence on the sympathetic nervous system in rats than has been observed in other species. For example, in rabbits, the sympathetic nervous system elevates nighttime IOP, but a secondary mechanism lowers daytime IOP [8, 52-54]. In mice, the IOP rhythm has been tied to local circadian oscillators, NE signals, and glucocorticoid/corticosterone inputs [44, 45, 51]. Thus, it appears that the rat rhythm is uniquely dependent upon sympathetic input.

### Alternative Explanations

One recent study relates pupil diameter to IOP demonstrating that dilation crowds the iridocorneal angle increasing outflow resistance thus elevating IOP [58]. Because the iris is under sympathetic control, one might attribute the IOP rhythm to a change in pupil size; however, both existing literature and the results presented herein argue against this. First, the rat IOP rhythm exists in constant darkness as shown in Fig 1A, and when the light conditions are altered the IOP rhythm re-entrains over several days whereas the pupil responds immediately to abrupt changes in ambient light including those at the transition to a different photoperiod [46, 59]. Further, while no studies have quantified the effects of pupil size on IOP directly in rats, one study in rabbits found that therapeutic dilation of the pupil elevated IOP by up to 5.2mmHg [60] despite the amplitude of the rabbit IOP rhythm being notably higher, ranging from 6.4 to 16.6mmHg [61], refuting the claim that the IOP rhythm is purely driven by pupil diameter.

Another argument might be that the bilateral SCGx performed in this study disrupted pineal gland innervation thereby abolishing the systemic melatonin rhythm [62-64]. This explanation is also inconsistent with the findings of this work. Topical TTX was as effective as SCGx in eliminating the nocturnal IOP rise, though its effects were transient, in contrast to the permanent loss of rhythm following SCGx. The rat IOP rhythm cannot be explained by systemic melatonin, as topical TTX produced only local ocular effects and thus would not have interfered with pineal melatonin production. Further, local melatonin production is unlikely to explain the IOP rhythm, as SCG regulates melatonin production in the pineal gland, but not in other tissues like the retina, which follows its own circadian clock [29-31].

The role of corneal sensory nerves was also considered as Aδ fibers in the cornea function as mechano-nociceptors and sense pressure changes which may potentially include IOP changes, though direct evidence of this is limited [65]. TTX is commonly used as a corneal anesthetic and would also affect this process. The presented SCGx results also eliminate the role of sensory neurons in the ocular surface as a driver of the IOP rhythm.

Finally, glucocorticoids have been implicated in the IOP rhythm of mice, raising the possibility of systemic hormonal cues as the driver of the IOP rhythm [10, 44]. Our results find that this is not the case with the rat IOP rhythm. The unilateral and transient suppression of nighttime IOP by TTX cannot be explained by systemic glucocorticoids, which would act also on the paired eye and persist beyond a single night [66]; the lack of lasting circadian changes further rules out disruption of hormonal rhythm as the driving mechanism.

### Future Work

Future studies should directly assess whether circadian clock genes in tissues interact with sympathetic efferents to modulate the IOP rhythm, as our current results argue against it but do not rule out such contributions. Additionally, future work should identify downstream processes through which efferent signals influence IOP to map the intermediate steps between neural input, ocular tissue response, and pressure regulation. Lastly, IOP changes are multifactorial and can be the result of any variation in the production-drainage relationship. As such, future work should elucidate which of these processes are under sympathetic control and to what degree they contribute to the circadian IOP rhythm.

## ACKNOWLEGDEMENTS

The work was supported by NIH grant R01 EY027037

